# Subtle left-right asymmetry of gene expression profiles in embryonic and foetal human brains

**DOI:** 10.1101/263111

**Authors:** Carolien G.F. de Kovel, Steven N. Lisgo, Simon E. Fisher, Clyde Francks

## Abstract

Left-right laterality is an important aspect of human brain organization for which the genetic basis is poorly understood. Using RNA sequencing data we contrasted gene expression in left- and right-sided samples from several structures of the anterior central nervous systems of post mortem human embryos and fetuses. While few individual genes stood out as significantly lateralized, most structures showed evidence of laterality of their overall transcriptomic profiles. These left-right differences showed overlap with age-dependent changes in expression, indicating lateralized maturation rates, but not consistently in left-right orientation over all structures. Brain asymmetry may therefore originate in multiple locations, or if there is a single origin, it is earlier than 5 weeks post conception, with structure-specific lateralized processes already underway by this age. This pattern is broadly consistent with the weak correlations reported between various aspects of adult brain laterality, such as language dominance and handedness.

## INTRODUCTION

The human brain shows left-right laterality of its anatomy and function ^1-8^. For example, more than 85% of people have left-hemisphere language dominance ^9^, and roughly 90% of people are righthanded ^10^. Possible reasons for these brain asymmetries are still debated, but various cognitive disorders can involve deviations from typical asymmetry patterns, suggesting that laterality is an important aspect of optimal brain development^11-13^

Anatomical laterality of various brain structures becomes apparent *in utero* ^14–19^, with the earliest observation at just 11 post conception weeks (pcw), concerning the choroid plexuses (left larger than right on average^14^). In addition, lateralized arm movements have been observed as early as 8 pcw ^20,21^. In a previous study, we showed that the spinal cords and hindbrains of human embryos aged in the range of 4 to 8 pcw showed left-right differences in gene expression that were related to a difference in maturation rates of the two sides^22^. However, we did not have gene expression data on more anterior regions of the developing CNS^22^, while published studies on cerebral cortical laterality of gene expression had only been performed on older foetal samples, ranging from 12 to 25 pcw ^23-25^. One of these studies found evidence for laterality of expression of the transcription factor LMO4 and various other genes, and also suggested a developmental asynchrony between hemispheres^23^.

In order to assess transcriptomic laterality in anterior CNS regions at earlier stages than previously studied, here we generated a new RNA sequencing dataset based on the left and right forebrains and midbrains of human embryos aged 5-5.5 pcw, for which healthy pregnancies had been terminated by voluntary medical abortions. In addition, the Human Developmental Biology Resource (UK) recently released a transcriptomic dataset in which various structures of the developing brain, in the age range 7.5-14 pcw, had been separated into left and right prior to RNA sequencing ^26^, which included cerebral cortex separated into temporal and non-temporal lobe, basal ganglion, diencephalon, and choroid plexus of the lateral ventricles. No laterality-related analysis of these data had been performed prior to that which we report here. A third publicly available dataset from embryos aged 4.5-9 pcw included various structures which had not been separated into left and right ^26^, but was nonetheless useful in assessing age-dependent changes of gene expression spanning the range of 5.5.5pcw.

## RESULTS AND DISCUSSION

Note: detailed structure-by-structure results are available in Supplementary Information.

### Developmental asynchrony is a general feature across most fore- and midbrain structures

Post-quality-control sample numbers for each structure and age range are listed in Table 1. Many genes increased or decreased their expression bilaterally with embryonic/foetal age. There was a general pattern of transitioning from gene expression profiles of cellular proliferation to neuronal differentiation with increased age (Supp. Table 3). This pattern was similar to the spinal cord and hindbrain at ages 4-8pcw, as we previously described ^22^, and is consistent with the normal development of neural tissue ^27,28^. In most of the structures we saw correlations between the effects of age and the effects of side on gene expression levels, with the exception of the embryonic forebrain at 5-5.5pcw, in which laterality of expression was not obviously linked to age-dependent changes (Fig. 1, Fig. 2, Fig. 3). We interpret such correlations as subtle differences in maturation rates between the left and right sides of a given structure (Fig. 2, Fig. 3). In other words, one side typically led the other slightly in terms of the developmental changes which both sides followed. Cell birth timing analysis in zebrafish has also shown an asynchrony of neurogenesis between the left and right habenula, where the timing influences the types of differentiated neurons that develop subsequently ^29^.

**Table 1.** Overview of included samples with left/right data

| ID | Age (pcw) | Sex | Source | BG* | CC* | CP* | DE* | TC* | FB* | MB* |
| --- | --- | --- | --- | --- | --- | --- | --- | --- | --- | --- |
| S13128 | 5 | Male | This study |  |  |  |  |  | Y | Y |
| S13048 | 5.5 | Female | This study |  |  |  |  |  | Y | Y |
| S13052 | 5.5 | Male | This study |  |  |  |  |  | Y | Y |
| S13097 | 5.5 | Male | This study |  |  |  |  |  | Y |  |
| S13192 | 5.5 | Male | This study |  |  |  |  |  |  | Y |
| S13290 | 5.5 | Male | This study |  |  |  |  |  | Y | Y |
| 11831 | 7.5 | Male | Lindsay *et al.* 2016 | Y | Y | Y | Y |  |  |  |
| 11823 | 8 | Male | Lindsay *et al.* 2016 |  |  | Y | Y |  |  |  |
| 11826 | 8 | Female | Lindsay *et al.* 2016 |  | Y |  | Y |  |  |  |
| 11827 | 8 | Female | Lindsay *et al.* 2016 | Y | Y |  | Y |  |  |  |
| 11846 | 8 | Female | Lindsay *et al.* 2016 | Y | Y | Y | Y |  |  |  |
| 11918 | 8 | Female | Lindsay *et al.* 2016 | Y | Y |  |  |  |  |  |
| 11920 | 8 | Male | Lindsay *et al.* 2016 | Y |  |  |  |  |  |  |
| 11830 | 8.5 | Female | Lindsay *et al.* 2016 |  |  |  | Y |  |  |  |
| 11832 | 8.5 | Female | Lindsay *et al.* 2016 | Y |  |  | Y |  |  |  |
| 11820 | 9 | Female | Lindsay *et al.* 2016 |  | Y | Y |  | Y |  |  |
| 11845 | 9 | Male | Lindsay *et al.* 2016 |  | Y | Y | Y | Y |  |  |
| 11851 | 9 | Female | Lindsay *et al.* 2016 | Y |  |  |  |  |  |  |
| 11873 | 9 | Female | Lindsay *et al.* 2016 | Y |  |  | Y |  |  |  |
| 11841 | 10 | Male | Lindsay *et al.* 2016 |  |  |  | Y |  |  |  |
| 11833 | 11 | Female | Lindsay *et al.* 2016 | Y |  | Y | Y |  |  |  |
| 11930 | 11 | Female | Lindsay *et al.* 2016 |  | Y |  |  |  |  |  |
| 11942 | 11 | Female | Lindsay *et al.* 2016 | Y | Y |  |  | Y |  |  |
| 11834 | 12 | Male | Lindsay *et al.* 2016 |  |  | Y |  | Y |  |  |
| 11885 | 12 | Male | Lindsay *et al.* 2016 |  | Y |  |  | Y |  |  |
| 12007 | 12 | Male | Lindsay *et al.* 2016 | Y | Y | Y | Y |  |  |  |
| 1923 | 13 | Female | Lindsay *et al.* 2016 | Y | Y |  | Y |  |  |  |

\* BG = basal ganglia, CC = cerebral minus temporal cortex, CP = choroid plexus of the lateral ventricles, DE = diencephalon,
TC = temporal cortex, FB = forebrain, MB = midbrain.

**Fig. 1.**
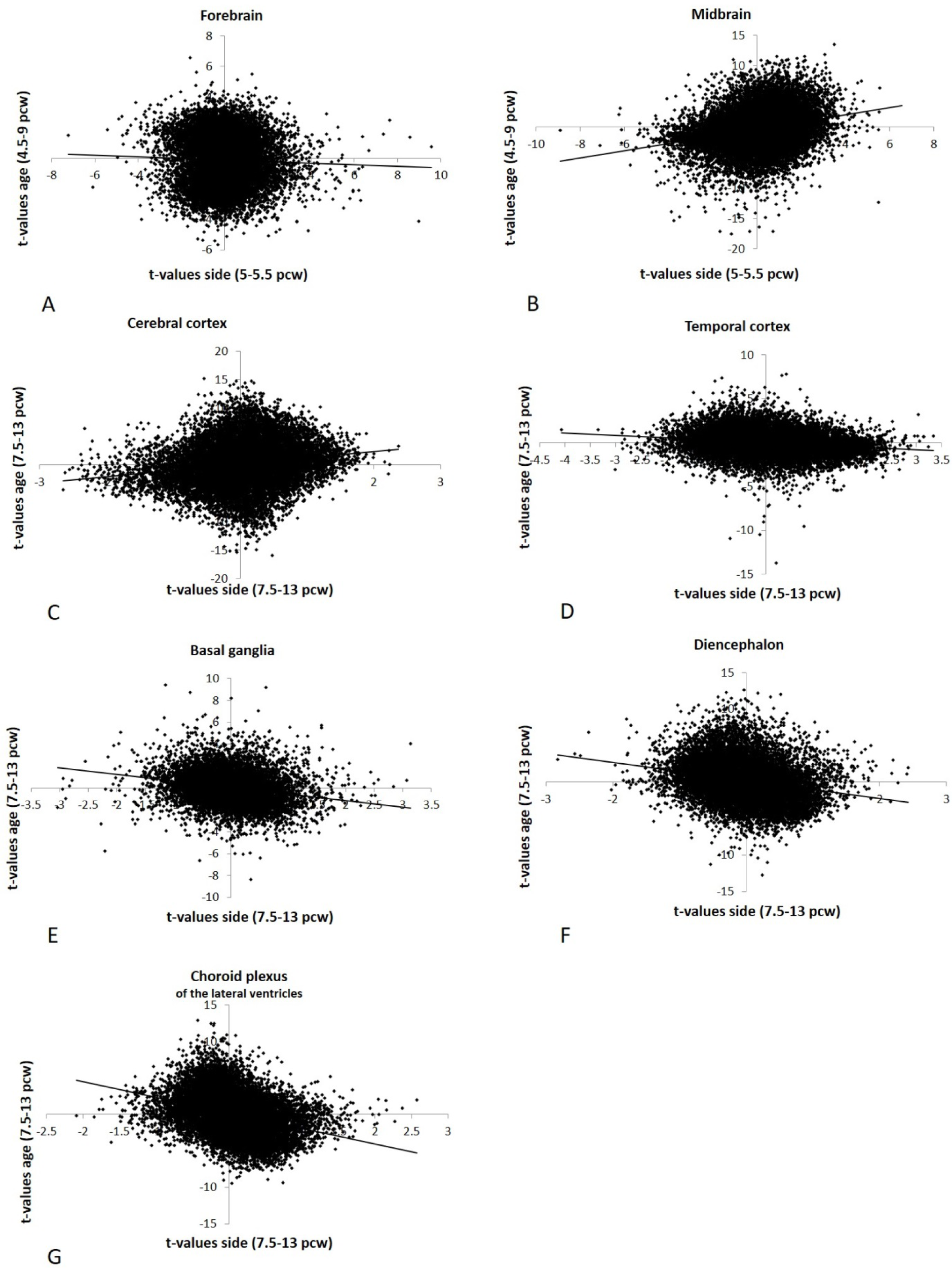
Relation between effects of age and side on gene expression (as t-values) for the brain structures aged 5,5 pcw and aged 7.5-13pcw. Each dot is an individual gene. X-axis: left-right differential expression t value, with positive values indicating genes with higher right-sided expression. Y-axis: age-effect t value, where positive t values indicate genes which increase in expression with age. A positive age-side correlation therefore indicates that the right side of a structure leads the left side in the transcriptional changes which both sides follow. R- and p-values can be found in the main text. Note that for the midbrain and the forebrain t-values for age and side were not computed on the same samples (see main text).

**Fig. 2.**
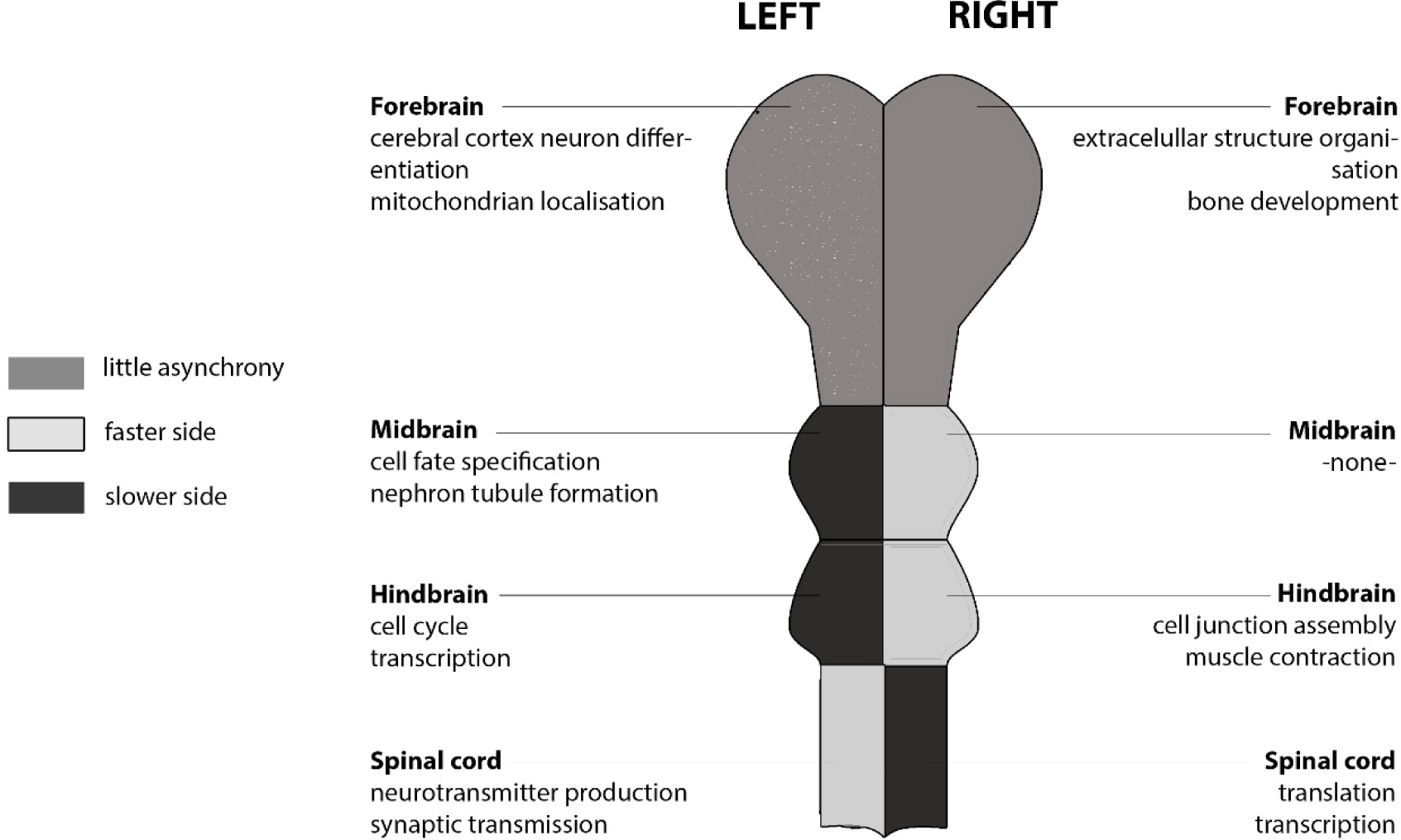
Schematic representation of the embryonic central nervous system around age 5-5.5 pcw. The lighter shade of grey indicates the side of the structure that has a higher expression of genes that increase in expression with age (the ‘faster’ side). Two top processes from the GO-enrichment analysis are shown for each side of each structure. Multiple GO-terms have sometimes been collapsed into a single process. Spinal cord and hindbrain were previously described in De Kovel et al. (2017).

**Fig. 3.**
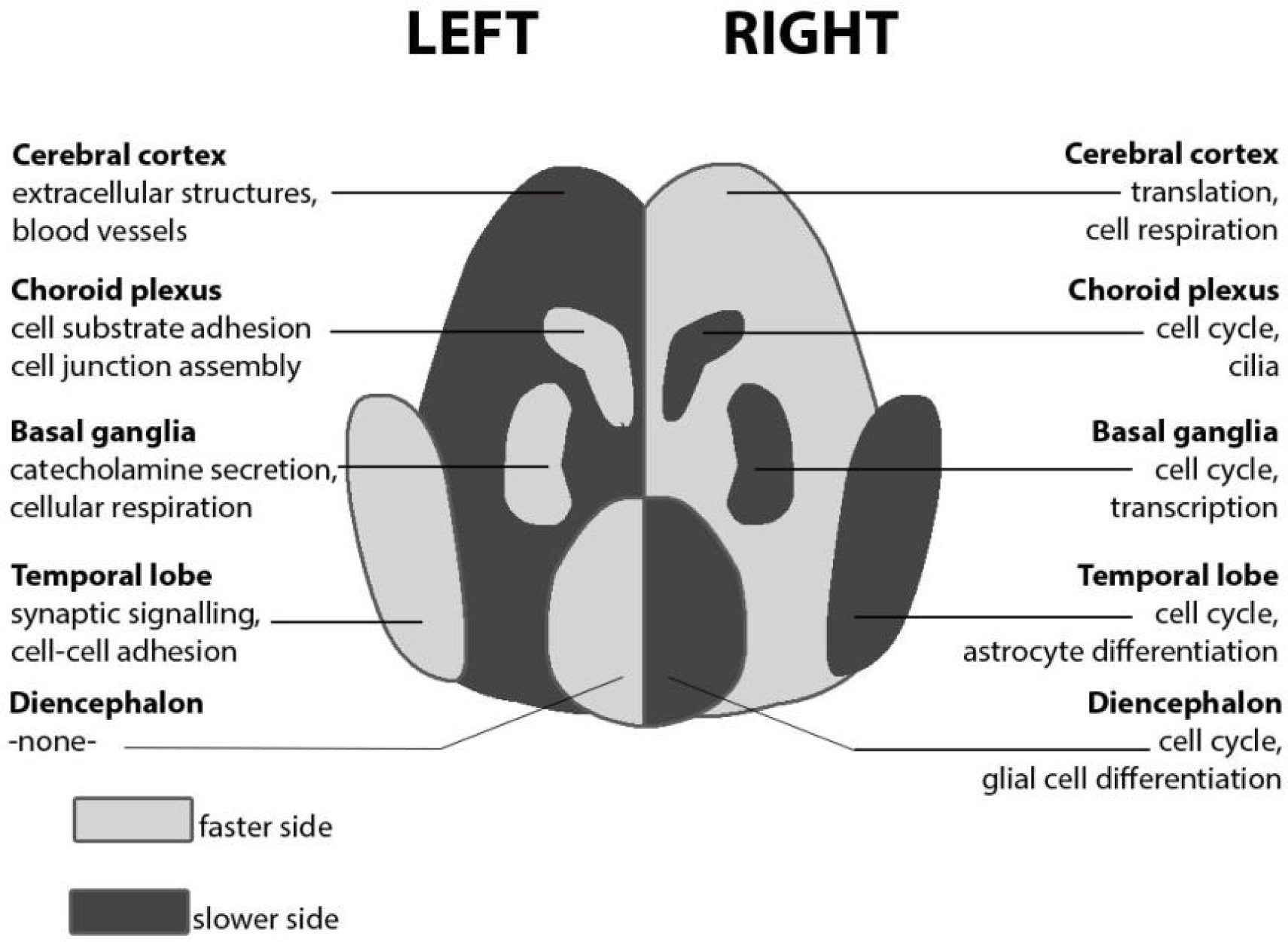
Schematic representation of anterior sub-structures of the foetal brain at 7.5-13 pcw. The lighter shade indicates the side of the structure that has a higher expression of genes that also increase in expression with age (i.e. the ‘faster’ side). For each side and each structure, the top two biological processes showing higher expression than the contralateral side are shown. (Multiple GO-terms have sometimes been collapsed into a single process for illustrative purposes.)

Table 1 Overview of included samples with left/right data

### Structure-specific orientations of developmental asynchrony

In a previous study, we reported faster maturation of the left spinal cord relative to the right in 4-8pcw human embryos, with the opposite pattern in hindbrain, which we attributed potentially to the crossing-over of major nerve tracts from spinal cord to hindbrain ^22^ (Fig. 2). One expectation going into the present study was therefore that, since most nerve fibre tracts pass ipsilaterally from hindbrain to more anterior regions, the midbrain and forebrain structures would consistently show the same developmental asynchrony as the hindbrain. This was borne out in the current study, insofar as the midbrain at 5-5.5pcw showed developmental asynchrony similar to hindbrain, i.e. the right side ahead, while the forebrain at this stage showed barely any difference in maturation rates between the two sides (Fig. 1, Fig. 2). Another recent study of the spinal cords of older human embryos, aged 8-12pcw, also found gene expression left-right differences that were broadly consistent with those we reported, as well as methylation differences ^30^.

However, in the present study, there was a very weak side-age relation in the forebrain at 5-5.5pcw (side-age correlation R=-0.04) which was opposite in direction to that in the midbrain at this stage, and there was also significant laterality at the level of functional gene sets in the forebrain at 5-5.5pcw which was not obviously related to age-dependent changes occurring from 4.5-9pcw (see further below). In addition, within the older age range 7.5-13pcw, the left side matured faster than the right for the temporal lobe, diencephalon, basal ganglia and choroid plexus, and only the cerebral cortex minus temporal lobe showed faster right-side transcriptional development. Therefore, taking together the current study and our previous study of spinal cord and hindbrain, it appears that diverse regions of the developing CNS have already embarked on subtly asynchronous programs of development by 5pcw, without an overall pattern at that stage, or subsequently, that can be explained solely in terms of the brain to spinal cord crossover (Fig. 2, Fig. 3).

Our previous study of the spinal cord and hindbrain was motivated by observations of embryonic laterality of arm movements at 8pcw ^20^, i.e. at a stage prior to the innervation of the descending corticospinal tracts into the spinal cord ^31^, so that the lateralized movements are probably not under forebrain control. We suggested that the spinal cord may be a developmental origin of asymmetry which could even cause later asymmetry in the cerebral cortex, via the formation of lateralized connectivity ^22^. This remains possible, at least in part, but the current study shows transcriptional laterality of the forebrain already at 5pcw, before corticospinal and spinocortical neural tracts are established.

The observation that laterality is already proceeding in multiple structures during embryonic and foetal development, without a simple and overall pattern to one side or the other, is consistent with the fact that individual differences in diverse aspects of adult brain laterality are poorly correlated with each other. For example, although the large majority of humans have left-hemisphere language dominance and right-handedness, most left-handed individuals also have left-hemisphere language dominance ^9,32^. With regard to motor behaviour specifically, it may be that the spinal cord is a developmental origin of laterality which then affects later cerebral cortical laterality for handedness, but not necessarily for other cerebral cortical functions and regions.

### Shared and structure-specific laterality of biological processes

From 7.5-13pcw, the basal ganglia, temporal cortex, diencephalon, and choroid plexus showed increased right-sided expression of processes involved in mitosis (a transcriptomic reflection of cellular proliferation), sometimes accompanied by higher expression of the target genes of E2F transcription factors, as was also observed previously for right spinal cord ^22,30^ (Supp. Table. 1 and 2). Overlaps between lateralised GO-terms in the various structures, as shown by the Jaccard index, were clearly larger in the right sides of the various structures, excluding cerebral cortex, than in the left sides (Supp. Fig. 7). There were also structure-specific lateralities of biological processes which we discuss in this section:

Asymmetric development of the cerebral cortex is probably the most obvious anatomical asymmetry of the developing foetal brain by mid gestation ^17,19^. Various studies have shown that while the left hemisphere is larger in the perinatal period ^17,18^, the right hemisphere seems to mature faster functionally ^33-36^. The most significant left-right differences in the cerebral minus temporal cortex involved gene-sets for angiogenesis and extracellular structure organisation (Fig. 3, Supp. Table 1), which were more strongly expressed on the left. Extracellular matrix molecules are involved in such diverse processes as neural stem cell differentiation, neuronal migration, the formation of axonal tracts, and the maturation and function of synapses in the central nervous system matrix ^37^. Blood vessels can play many roles in CNS tissue, including guiding axon outgrowth and neuronal migration, in addition to providing the developing tissue with oxygen ^38^.

The forebrain at 5-5.5pcw, which goes on to differentiate into cerebral cortex, basal ganglia, and diencephalon at later stages, showed higher expression of genes involved in angiogenesis (blood-vessel formation) on the right (Supp. Table 1), i.e. the opposite laterality to the cerebral minus temporal cortex at 7.5-13pcw. Possibly this is a transient process, first upregulated broadly on the left, but then becoming more region-specific in its later laterality.

Other notable asymmetries involved gene sets in the basal ganglia which affect dopamine secretion, as well as the targets of the Androgen Receptor (AR) transcription factor in the diencephalon (Supp. Table 2). Dopamine is very important in the functioning of the basal ganglia ^39^, while steroid hormone pathways have long been studied in relation to variability in brain anatomical laterality ^40,41^, handedness ^42,43^, and language-related development ^44,45^. The transcriptional asymmetries which we observed in these developing subcortical structures may therefore play important roles in creating broader functional lateralities for motor and language functions, also involving the cerebral cortex.

The choroid plexus is of particular interest, given that it shows the earliest reported anatomical laterality in the developing human brain, at 11pcw ^14^, and has been hypothesised to affect broader asymmetries through lateralized control of cerebrospinal fluid in the ventricles ^46^. The choroid plexuses were clearly distinct from the other brain structures in the age range 7.5-13pcw in terms of their overall transcriptomic profile (Supp. Fig. 3, Supp. Table 5), being distinguished by relatively high expression of genes involved in the extracellular matrix, cell adhesion, immune responses and ciliary function (Supp. Table 5, Fig. 3). In our analysis, genes involved in cell adhesion and immune responses were more highly expressed in the left choroid plexus, and cilia-related gene-sets on the right (Supp. Table 1). In contrast to other parts of the CNS, immune responses are found in healthy tissues at the blood-brain barrier and the blood-cerebrospinal fluid (CSF) barrier, such as the choroid plexus ^47,48^.

The expression of genes involved in ciliary function is consistent with the role of the choroid plexus in circulating cerebrospinal fluid in the ventricles ^49^. Cilia, with their unidirectional rotation which derives ultimately from the molecular chirality of amino acids, have also been shown to play an important role in establishing the left-right body axis, particularly within the embryonic node (a pitted structure located at the ventral embryonic surface) ^50-52^. Whether cilia play a similar role in central nervous system asymmetry is not clear. People with a complete reversal of the left/right pattern of the visceral organs in Kartagener syndrome, a primary ciliary dyskinesia (MIM 244400), have shown the same proportions of cerebral dominance for handedness and auditory language dominance as the general population ^53,54^. Nonetheless, the high expression of cilia-related genes in the choroid plexus, and its early anatomical laterality in humans, makes it a plausible candidate as a developmental origin of laterality in the CNS.

### Putative genetic links to visceral left/right patterning and zebrafish brain left/right patterning

Asymmetries of the brain and/or behaviour have also been observed extensively in other vertebrates ^55-59^ and invertebrates ^60-62^. Some molecular mechanisms which underlie developmental asymmetries of the nervous system have begun to be described in model species such as *C. elegans* ^63^, *Drosophila* ^64^, and particularly in the zebrafish ^65^. In addition, various organs of the vertebrate viscera (heart, lungs etc.) show asymmetries in terms of size, shape or positioning ^66^. The developmental origins of left/right visceral patterning can be observed shortly after gastrulation ^67,68^, and many genes have been identified which are involved in this patterning.

In terms of individual genes, we found *KCTD12* and *SNAI1* to be asymmetrically expressed towards the right in the 5-5.5pcw forebrain after false discovery rate correction, as well as targets of the transcription factor FOXJ1. We note, however, that these individual gene findings would not survive further statistical correction for multiple testing over all the different structures analyzed in this study; nonetheless it is notable that these were the top genes in our analysis, for the following reasons:

The fish homologue of *KCTD12 (Kctd12.1* or *lov*) is involved in left-right asymmetrical development of the fish brain, and is asymmetrically expressed towards the left in zebrafish diencephalon during the phase ~40-96 hours post fertilisation ^69,70^. In zebrafish, the parapineal organ of the embryonic dorsal diencephalon migrates to the left side, influenced by the bilaterally expressed protein Fgf8 ^71^. The habenulae, which at this early stage are clusters of cells located to the left and right of the pineal gland, then go on to develop structural and functional asymmetries. These involve left-right differences in the proportions of neuronal subtypes with distinct patterns of gene expression, axon terminal morphology and connectivity ^69^. An important factor is asymmetric nodal (*Ndr2*) signalling, and later asymmetric expression of *Kctd12.1 (lov), Kctd8 (dex), Pitx2*, and other genes.

Both *SNAH* ^*72*^ and *FOXJ1* ^73-75^ have also been shown to be involved in left-right visceral asymmetry. Furthermore, in the zebrafish mutant fsi (frequent *situs inversus*), structural asymmetries in the diencephalon are correlated with visceral asymmetries. Some behavioural asymmetries are also left-right reversed in mutant fish with reversed visceral asymmetry, while other behaviours retain the wildtype direction ^76^. In humans the relation between visceral and brain asymmetry is also complex. As noted above, people with a complete reversal of the left/right pattern of the visceral organs in Kartagener syndrome (a primary ciliary dyskinesia) have shown the same proportions of cerebral dominance for handedness and auditory language dominance as the general population ^53,54^, except when the sample size was extremely small (3 patients)^77^. However, some structural asymmetries of the brain may be reversed in people with reversed visceral laterality, although again the sample sizes in these studies have been extremely small ^78,79^. One study indicated that genes involved in visceral laterality are enriched for associations of common polymorphisms with a measure of human handedness, although the sample size was relatively small for a study of a weakly heritable and complex trait ^80^.

### General conclusions

In this study, starting from 5 weeks post conception, the left and right sides of developing human brain structures showed subtle differences in their global profiles of gene expression. Various structures showed transcriptomic left-right asynchrony, although not consistently in orientation over all structures. This suggests that different aspects of adult human brain laterality are likely to have relatively distinct developmental histories going as far back as the embryo, and sheds new light on the timing of these lateralized programs of development. While many of the structures had higher expression of processes involved in cell proliferation on their ‘slower’ sides, there were also lateralities involving biological processes that are relatively structure-specific. Furthermore, our data did not indicate any one structure as a unique developmental origin of laterality in the human CNS, in terms of preceding the others in lateralized development. Studies earlier in development than 4-5pcw would be required to resolve whether the brain laterality is triggered by only one mechanism, or a number of different structure-specific mechanisms.

Transcriptomic studies often form the basis of subsequent gene-functional studies, which are typically based on selecting a small number of individual genes for detailed biological analysis. However, such a reductive approach may not be well motivated by our results, given that the signals in our data were primarily at the global transcriptomic (i.e. the age-side relations) or multi-gene (gene set) levels. One promising avenue for follow-up may be to try observing anatomically localized left-right asymmetries of cell type abundancies, although avoiding a confound of left-right with depth in e.g. slide microscopy would be challenging. There remains a bridge to cross from genomic-level analysis to individual gene functional analysis, but the genomic level is itself an informative level to probe and report, and reflects well the complexity and diversity of processes already showing laterality by five weeks post conception and thereafter.

### Limitations

The evidence for laterality at the level of individual genes was much weaker than for gene-sets, with only the midbrain and forebrain at 5-5.5pcw showing small numbers of significantly lateralized individual genes after false discovery rate correction, which would not be significant after further correction for testing all structures in this study. Studies of gene expression laterality in post mortem human embryos and fetuses are necessarily limited to relatively small numbers of samples ^22,23,28,30^ As the left and right sides of the central nervous system are grossly anatomically similar, any left-right differences of gene expression are expected to be subtle overall. When testing each individual gene in a transcriptomic dataset and performing appropriate statistical correction for multiple testing over thousands of genes in small samples, the power is low to detect subtle laterality at the level of individual genes. Transcriptomic-level and gene-set-based analyses have a greater sensitivity relative to individual gene testing, for detecting subtle gene expression differences between closely matched samples, as has been reported before ^81,82^.

Another possible limitation is the accuracy of the dissections. A bias in the separation of left and right structures at the midline could potentially influence the results, if such a bias was consistent across embryos/foetuses. For example, the left-right axis might become conflated with anterior-posterior or dorsal-ventral gene expression gradients. The anatomists at HDBR are highly skilled in dissecting embryonic and foetal human brains, but the possibility cannot be totally excluded.

A third limitation regards the spatial resolution of the datasets. Clearly each of the structures analysed has a complex internal composition of diverse cell types, sub-tissues and regions. For example, different zones of the foetal cerebral cortex may be differentiating from each other in terms of their gene expression ^26^, so that the left-right differences we observed will have been a global average over different sub-regions. Consistent with this, we observed contrasting patterns of the temporal cortex versus the rest of the cerebral cortex (Fig. 3), which was the finest resolution of the cerebral cortex possible in our study.

## MATERIALS & METHODS

### Embryos at 5 to 5.5 weeks post conception

#### Collection, library _preparation and sequencing

The data for embryos aged Carnegie stages CS15-CS16 were generated newly for this study. Six embryos were collected by the MRC/Wellcome-Trust funded Human Developmental Biology Resource (HDBR) (United Kingdom) of CS15 and CS16, therefore estimated between 5 and 5.5 weeks after conception. The embryos were obtained anonymously from voluntary medical terminations (a combination of mifepristone and misoprostol) or physical termination following appropriate informed consent by the donors, and with ethical approval from the Newcastle and North Tyneside NHS Health Authority Joint Ethics Committee. Donors to HDBR are asked to give written consent for the embryonic material to be collected, and are only approached once a decision to terminate their pregnancy has been made. The development of the embryos was assessed and designated to the relevant Carnegie stage (CS) ^83^, using a practical staging guide devised to enable staging to a particular CS and using the external morphology of a single sample ^84^. Forebrain and midbrain were separated and then dissected into left and right sides. RNA was extracted, subjected to quality control, and then shipped to the genomics service provider Beijing Genomics Institute (BGI) (Shenzhen, China; www.genomics.cn). A detailed description of the procedure can be found in ^22^. The embryos were five males and one female. The female was from a physical pregnancy termination, the males from chemical pregnancy terminations.

Oligo(T)’s were used to enrich for mRNA out of the pool of total RNA. After fragmentation, cDNA was generated with random hexamer primers, and the cDNA fragments were connected with adapters, following standard procedures ^85^. Barcoded cDNA fragments were sequenced in two lanes of an Illumina HiSeq 4000 sequencer. We used paired-end sequencing with a read length of 100 bases. Raw reads were filtered to exclude reads with more than 5% unknown bases, reads which contained more than 20% bases with quality score below 15, and reads with adapters. After filtering, the median size was 5.65 Gb per library (range 3.66 - 7.26 Gb).

#### Data processing

RNA sequencing data were produced as fastq-files. FastQC (v0.11.5, Babraham Bioinformatics, Cambridge, UK) showed that the quality was very good without overrepresented sequences or adapters, and phred base quality score mostly > 37. GC content varied between 48% and 50%. Reads were aligned to the Human reference GRCh38 from UCSC (http://genome-euro.ucsc.edu) using Hisat2 (v2.0.4, ^86^). Using the same reference with RefSeq gene annotations, reads were counted per gene using RSEM (v1.3.0 ^87^). Both packages use bowtie2 ^88^.

#### Further quality control

A separate pipeline using HiSat (v0.1.6 beta, for mapping against hg19) and GATK (v3.4.0 ^89^) was used to create genotype calls for single nucleotide polymorphisms (SNPs) from the RNA sequencing data. The SNP data were used in plink (v1.07 ^90^) to confirm that left and right pairs of matched samples came from the same individual, and to confirm the sexes of the samples. A second confirmation for sex was found by looking at the expression data for the X-chromosomal gene *XIST* and the Y-chromosomal genes *EIF1AY* and *KDM5D* (See Supp. Table 6). Clustering analysis on expression data in R showed that forebrain and midbrain separated into two clusters. The right side of one of the forebrain samples fell into the midbrain cluster. Data for the forebrain were discarded for this embryo. In addition, both sides of one midbrain sample clustered in the forebrain group. Data for the midbrain were discarded for this embryo. The discarded midbrain and forebrain data were not from the same embryo. Therefore, after this extra quality control procedure, we had data from four males and one female for each structure, though not exactly the same embryos for hindbrain and midbrain (See Table I for an overview of the samples). In R (version 3.3.2), expression data were normalized and transformed into log2 cpm (counts per million). Genes were filtered to retain only those for which at least three libraries had at least five reads per gene, separately for each brain structure.

### Foetuses aged 7.5 to 13 weeks post conception

#### Collection, library preparation and sequencing

Data for foetuses aged 7.5-14pcw were made publicly available by the HDBR as described previously ^26^, but quality control steps for the present study (see below) meant that we were only able to analyse the age range 7.5-13pcw. To our knowledge, no analyses of these data had been performed in relation to laterality, i.e. no comparisons of left and right gene expression were published from this dataset, prior to the present study. Ages younger than 9 PCW in this dataset were estimated from anatomical-based Carnegie staging, but for simplicity we refer to the whole dataset in PCW units, as did Lindsay et al. 2016. Collection and dissection of tissue, RNA isolation and sequencing, and SNP genotyping, were described previously ^26^. In brief, tissue was collected from foetuses after elective termination of pregnancy. Brains were dissected, with the level of subdivision depending on the age and size of the foetus. Tissues were sent to AROS Applied Biotechnology (Aarhus, Denmark; www.arosab.com), who prepared DNA and RNA and carried out SNP genotyping and RNA-sequencing. After generation of cDNA, libraries were generated with TruSeq Stranded mRNA LT sample preparation kit (Illumina, San Diego, CA). The libraries were sequenced on an Illumina HiSeq2000. Sequencing was paired-end, with a read length of 100 bases. For the present study, we downloaded Fastq-files, containing the results of the RNA-sequencing, from E-MTAB-4840 in the ArrayExpress database (http://www.ebi.ac.uk/arrayexpress/experiments/E-MTAB-4840/), along with the meta-data and, when available, SNP genotype data (E-MTAB-4843).

#### RNA-sequencing library selection

The entire dataset included RNA-sequencing libraries with annotations to various structures of the developing central nervous system, but as described below, sample size and quality control considerations meant that we were ultimately only able to analyse data from libraries annotated as ‘cerebral cortex’, ‘temporal lobe’, ‘basal ganglion’, ‘diencephalon’ and ‘choroid plexus’. As described previously ^26^, the temporal lobe had been dissected from the rest of the cerebral cortex.

Since we were interested in laterality of gene expression, we only used data for a given structure when a separate library was available for both the left-and-right matched pair of samples from a given foetus. One foetus was excluded for the temporal cortex, because both libraries were annotated as ‘right’, which could not be resolved. Additionally we excluded the ‘forebrain’ and ‘telencephalon’ libraries since these structures had been divided into sub-structures for many of the foetuses, and we were interested in the highest resolution analysis possible (i.e. ‘forebrain’ includes all cerebral cortical regions, basal ganglia and diencephalon, while ‘telencephalon’ includes all cerebral cortical regions and basal ganglia). The choroid plexus annotations corresponded specifically to the cerebral choroid plexus, i.e. not including the choroid plexus of the third and fourth ventricles (personal communication Suzanne Lindsay, 2017).

The ‘cerebral cortex’ samples and the ‘temporal cortex’ of larger brains had further been dissected into slices, with the most anterior slice labelled number 1 within each structure. For the cerebral minus temporal cortex there were usually five slices, for temporal cortex two slices. To avoid overrepresentation of foetuses with multiple slices per structure in our analyses, we selected one slice per structure per foetus. We chose the most anterior slice for which both left and right libraries were available. Three foetuses were not included for the cerebral minus temporal cortex, because no matching pair of slices was available.

#### Processing

We ran FastQC (v0.11.5, Babraham Bioinformatics, Cambridge, UK) on all fastq-files to check the quality. The results indicated that in many files, the Truseq adapters had not been completely removed. Also, the last two bases of a read never contained ‘A’. We therefore used Cutadapt (v1.11, ^91^) to remove the adapters according to the authors’ instruction, and to remove the last two bases. A new round of FastQC showed that we had succeeded. Median GC% was around 46% (range 37-48%).

Reads were aligned to the Human reference GRCh38 from UCSC using Hisat2 (v2.0.4, ^86^). Using the same reference with RefSeq gene annotations, reads were counted per gene using RSEM (v1.3.0 ^87^). Both packages used bowtie2 ^88^. This latter step was identical to the processing of the data on embryos aged 5-5.5pcw described above. Bcftools (v 1.3.1-173, ^92^) was used, with the same reference genome, to extract SNP calls from the RNA sequencing data. For most of the foetuses, SNP genotyping data had also been generated using Illumina’s HumanOmni5-Quad BeadChip (Illumina, San Diego, CA). We used Genomestudio 2.0 (Illumina (R)) to call the genotypes from the BeadChip data.

#### Further quality control

With plink (v1.07 ^90^), dimensions of multidimensional scaling (MDS) for the genotyped SNPs were plotted. Comparison with publicly available SNP-data indicated that the foetuses showed an ethnic distribution as could be expected from sample collection in the UK: the bulk of the samples coincided with European-descent origin, with tails towards both African and Asian descent (Supp. Fig. 4). SNP data were also used to confirm sexes. A second confirmation of sex was found by looking at the expression data for the X-chromosomal gene *XIST* and the Y-chromosomal genes *EIF1AY* and *KDM5D. XIST* expression should be much higher in females than in males, and expression of Y-chromosomal genes should be absent in females (See Supp. Table 6). All sexes could be confirmed. The SNPs within the exome, as extracted from the RNA sequencing data, were used to confirm that various structures truly matched which were assigned to the same foetus. All matched correctly. Some libraries in diencephalon had been split for sequencing. We added up the gene counts from these libraries for a given side.

In R (R version 3.3.2), we generated cluster plots, MDS-plots and heatplots to check for outliers in the expression data. One foetus was excluded for the cerebral cortex because it was a visible outlier. One foetus was excluded for choroid plexus because the left library was a visible outlier in the MDS-plot. One foetus was excluded for diencephalon because the left library was a visible outlier.

We set a minimum sample size threshold of five foetuses per structure to help ensure reliability of subsequent left-right differential expression analysis. Since the libraries from the oldest foetus had been discarded during quality control, the age range was now 7.5-13 pcw. For the included libraries, the median read count was 46.5 E06 (s.d. 7.7 E06, range 16.4 E06 - 64.4 E06). An overview of all included brain structures after quality control can be found in Table 1. In R (version 3.3.2), expression data were normalized and transformed into log2 cpm (counts per million). Genes were filtered to retain only those for which at least three libraries had at least five reads per gene, separately for each brain structure.

### Embryos aged 4.5 to 9 weeks post conception without left-right separation

Due to the narrow age range of 5-5.5 pcw for our newly generated fore- and midbrain left-right dataset, we could not test directly for age-related changes within this dataset itself. However, we also downloaded midbrain and forebrain data from the HDBR public dataset E-MTAB-4840 for embryos aged 4.5-9pcw which had not been dissected into left and right (and were therefore not part of the data described in the section above) ^26^. These data would be useful to assess bilateral age-dependent changes of gene expression (see below), over a developmental period which includes 5-5.5pcw. For forebrain, there were such data from 9 embryos (7 females, 2 males). For midbrain there were such data from 31 embryos (19 females, 12 males). These data were processed as described above.

### Data analysis

Once gene counts had been produced for the left-right data from the embryos aged 5-5.5pcw and foetuses aged 7.5-13pcw, further analysis was very similar for these two datasets, although they were analysed entirely separately. Any differences in the analysis are detailed below. In addition, the data on embryos aged 4.5-9pcw (not left-right separated) were treated similarly, with respect to bilateral differential gene expression by age (see below).

#### Differential expression analyses

Linear models with observational-level weights were fitted to obtain average expression values for each gene on each side (left or right), separately for each structure, and moderated t-statistics were used to assess differential expression between sides using the Bioconductor edgeR (version 3.6.8) and limma (version 3.22.7) packages with the *voom* option to provide shrinkage of variance for genes with low count numbers ^93-96^. Differential gene expression between the left and right sides of a structure was computed using a factor to distinguish individual embryos or foetuses, making this a paired analysis. Further covariates beyond this individual-level factor were not included. False discovery rate correction (Benjamini-Hochberg) was used to adjust for multiple testing within a structure.

Separately, to identify genes which were strongly affected by age or by sex, we also used models which included both age (a linear effect) and sex as factors, but then without the factor for distinguishing individual embryos or foetuses. In addition, certain of these analyses were performed across multiple structures (see below), in which case structure was added as a categorical factor.

#### Gene-set enrichment

Gene-set enrichment analyses can be used to test whether genes belonging to a particular functional set are over-represented in the results of, for example, a differential gene expression analysis. We used GSEA from the Broad Institute, Cambridge, UK (v2.0, build 45 ^97^) to test for gene-sets overrepresented among the genes which were expressed more highly within either the left or right side of a given structure, and also for genes that decreased or increased bilaterally in their expression with age. For these analyses, we first ranked genes according to their t-values in the left-right differential-expression analysis, and then used the ‘preranked’ option of the GSEA tool, with the option ‘weighted’ to take into account the t-values rather than the ranks. To define function-based gene-sets we used the gene ontology (GO) sets for ‘biological process’ (named C5 BP), which is a functional classification scheme for genes, as well as transcription factor (TF) targets sets (named C3 TFT), both as provided by the MSig database v5.2 ^97,98^. TFs are genes which control the spatiotemporal expression of other genes and are therefore crucial for regulated development. The target set of a given TF refers to all genes which have a sequence motif to which that TF could bind, located within 2000 bases from the gene’s transcription start site in the genome. Biological process or TF target gene-sets were only tested when at least 15 genes (and maximally 500 genes) belonging to the set were detected in the post-QC dataset for a given brain structure. The ‘preranked’ GSEA method does not use any cut-off on differential expression t-scores, but rather calculates an enrichment score that reflects the degree to which a set *S* is overrepresented at the extreme (either top or bottom) of the entire ranked list of detected genes. Normalisation corrects for the size of the gene-set leading to a normalised enrichment score (NES), and a family-wise error rate (FWER) is used to correct for multiple testing over the number of tested gene-sets in a given analysis ^97^.

When gene lists from differential expression analysis were compared to ‘all known genes’ rather than to a list of expressed genes in our datasets (as was appropriate for certain analyses: see below), PANTHER Overrepresentation Test (release 20160715), using GO Ontology database (released 201702-28) was used on the website http://geneontology.org/. Unless otherwise stated, the reference gene-sets used belonged to ‘GO biological process complete’.

To assess overlap of lateralised GO biological processes between the sides of the various brain structures, we calculated the Jaccard index ^99,100^ for GO terms that turned out to be enriched in the GSEA analyses with FDR < 0.05.

#### Permutations

We randomised the differential expression t-values with respect to the expressed genes for each separate left-right analysis, and ran GSEA for GO-terms again. This was repeated 10 times, and the number of gene-sets with FDR < 0.25 was noted. This was compared to the number for the true (nonrandomized) analysis of a given structure. If the number of gene-sets with FDR < 0.25 was orders of magnitude larger in the true observations than in the randomisations, this was interpreted as support for biologically meaningful left-right differences in the observed data, i.e. that groups of genes involved in the same biological processes were expressed more highly on one or other side of the tissue, rather than just random genes.

In addition, we performed permutations in which we randomly flipped, or left unflipped, the left and right samples of each embryo/fetus. The number of possible permutations depended on the number of embryos/fetuses in a given analysis (n) as 2^(n-1)^. For all permuted data-sets we carried out differential expression analyses, GSEA analyses and correlations between side effects and age effects on expression, as we did for the true observations. These permutations were done to check whether the left-right differences and patterns we found in the true data-set are an effect of consistent left-right differences in the brain structures, and not some artefact of accidental left-right differences at process-level. We plotted the r for correlations between side effects and age effects on expression against the similarity of the permuted dataset with the true observations (Supp. Fig. 5). We also recorded the number of GO-terms with FDR < 0.25 for each permuted dataset, and compared with the corresponding true data-set.

#### Further statistical testing

To identify MDS dimensions associated with sex, unpaired t-tests were used. For association of MDS dimensions with side (left/right), t-tests paired by individual embryo/foetus were used. Correlations of MDS dimensions with age were assessed by Pearson correlation. Multiple testing correction was done with the Bonferroni method for the number of dimensions relevant to a given analysis. To study correlation between the effects of age and side on the expression of genes, Pearson correlation between t-values from the age- and the side-differential expression analyses was computed. As the different genes will not be fully independent data points, the p-values for these correlations may be somewhat biased.

Note that for the descriptive analyses which we performed across structures, the structures with a larger number of available samples will have affected the analyses disproportionately. In addition, for most brain structures the data were not balanced for age and sex. If both age and sex influence asymmetric expression of genes, the unbalanced availability of data meant that these two effects on asymmetry will have been partly confounded, especially in the smaller sets. Given the small sample numbers, it was not possible to model these effects more thoroughly in left-right differential expression analysis, and instead we used a factor to distinguish individual embryos or foetuses to make a paired analysis, for which robust detection of laterality was the primary purpose. However, in the differential expression analysis by age, which was not paired, the measured effects of age could be imprecise due to confounding with sex, and we therefore did not attempt to calculate precise levels of the temporal asynchrony in development for the different structures when correlating side and age differential expression results. Nonetheless the overall age-dependent changes which we describe here are very much in line with the normal development of neural tissue (see Results and Discussion).

## Acknowledgements

We thank the women who donated the embryos and foetuses to the Human Developmental Biology Resource (UK). Thanks to Susan Lindsay for helpful input when writing the manuscript. Funding: C.G.F.de.K was supported by an Open Programme grant (824.14.005) to C.F. from the Netherlands Organization for Scientific Research (NWO). C.F. and S.E.F. are supported by the Max Planck Society (Germany).

## Author contributions

CGF de K: conceptualization, methodology, formal analysis, writing - original draft, writing – review & editing. CF: conceptualization, writing - original draft, writing - review & editing, project administration, funding acquisition. SNL: methodology, writing - review & editing. SEF: methodology, writing - review & editing.

## Additional information

The authors declare to have no competing or financial interests.

Supplementary information available online at ̤

Human embryos were obtained from the MRC/Wellcome-Trust funded Human Developmental Biology Resource (HDBR, http://www.hdbr.org, grant# 099175/Z/12/Z), with appropriate maternal written consent and approval from the Newcastle and North Tyneside NHS Health Authority National Research Ethics Committee (REC reference # 08/H0906/21+5). HDBR is licenced as a tissue bank by the UK Human Tissue Authority (HTA) and operates in accordance with all the relevant HTA Codes of Practice.

## Data availability

The following data set was generated: GSE99302, publically available at the NCBI Gene Expression Omnibus (http://www.ncbi.nlm.nih.gov/geo/).

The following previously published data sets were used: E-MTAB-4840, publically available from ArrayExpress (http://www.ebi.ac.uk/arrayexpress/experiments/E-MTAB-4840/ ^26^ and E-MTAB-4843, publically available from ArrayExpress (http://www.ebi.ac.uk/arrayexpress/experiments/E-MTAB-4843/ ^26^.

## Supplementary Files

Supplementary Information: detailed results and figures

Supplementary Table 1: Per structure GO-terms lateralised with FWER<0.05

Supplementary Table 2: Per structure Transcription Factor Target terms lateralised with FWER<0.05

Supplementary Table 3: Per structure GO-terms associated with age with FWER<0.05

Supplementary Table 4 GO-terms distinguishing forebrain from midbrain with FWER<0.05

Supplementary Table 5: Genes and GO-terms specific for choroid plexus

Supplementary Table 6: Genes differing in expression between males and females

